# *In vivo* ^18^F-DOPA PET imaging identifies a dopaminergic deficit in a rat model with a G51D α-synuclein mutation

**DOI:** 10.1101/2022.11.05.515268

**Authors:** Victoria Morley, Karamjit Singh Dolt, Carlos J. Alcaide-Corral, Tashfeen Walton, Christophe Lucatelli, Tomoji Mashimo, Adriana A. S. Tavares, Tilo Kunath

## Abstract

Parkinson’s disease (PD) is a neurodegenerative condition with several major hallmarks, including loss of *substantia nigra* neurons, reduction in striatal dopaminergic function, and formation of α-synuclein-rich Lewy bodies. Mutations in *SNCA*, encoding for α-synuclein, are a known cause of familial PD, and the G51D mutation causes a particularly aggressive form of the condition. CRISPR/Cas9 technology was used to introduce the G51D mutation into the endogenous rat *SNCA* gene. *SNCA*^G51D/+^ and *SNCA*^G51D/G51D^ rats were born in Mendelian ratios and did not exhibit any severe behavourial defects. *L*-3,4-dihydroxy-6-^18^F-fluorophenylalanine (^18^F-DOPA) positron emission tomography (PET) imaging was used to investigate this novel rat model. Wild-type (WT), *SNCA*^G51D/+^ and *SNCA*^G51D/G51D^ rats were characterised over the course of ageing (5, 11, and 16 months old) using ^18^F-DOPA PET imaging and kinetic modelling. We measured the influx rate constant (*K*_*i*_) and effective distribution volume ratio (*EDVR*) of ^18^F-DOPA in the striatum relative to the cerebellum in WT, *SNCA*^G51D/+^ and *SNCA*^G51D/G51D^ rats. A significant reduction in *EDVR* was observed in *SNCA*^G51D/G51D^ rats at 16 months of age indicative of increased dopamine turnover. Furthermore, we observed a significant asymmetry in EDVR between the left and right striatum in aged *SNCA*^G51D/G51D^ rats. The increased and asymmetric dopamine turnover observed in the striatum of aged *SNCA*^G51D/G51D^ rats is similar to prodromal PD, which suggests the presence of compensatory mechanisms. *SNCA*^G51D^ rats represent a novel genetic model of PD, and kinetic modelling of ^18^F-DOPA PET data has identified a highly relevant early disease phenotype.

## INTRODUCTION

Parkinson’s disease (PD) is a common progressive neurodegenerative condition affecting ∼1% of people over the age of 60 (Reeve et al., 2014). The condition results in deficits of motor function including abnormal posture and gait, tremor and rigidity (Gelb et al., 1999; Lees et al., 2009). Characteristic neuropathological findings include the loss of neurons from the *substantia nigra pars compacta* (SNpc) which project axons to the striatum where dopamine (DA) is released (Bernheimer et al., 1973). The pathological hallmark of PD is the formation of Lewy bodies and Lewy neurites, complex intracellular inclusions abundant in α-synuclein protein (Gibb and Lees, 1988; Spillantini et al., 1997). Cases of familial PD have been shown to result from mutations in *SNCA*, encoding α-synuclein, and the G51D mutation (c.152 G>A) has been identified to cause an early onset and aggressive form of PD with associated dementia and multiple system atrophy (Kiely et al., 2013; Lesage et al., 2013).

It has been extensively reported that PD patients have a decreased number of tyrosine hydroxylase (TH) positive neurons in the striatum when compared to controls, and this occurs when the condition has significantly progressed (Huot et al., 2007; Kordower et al., 2013). However, prior to significant neuronal loss, earlier deficits in presynaptic dopaminergic function in the striatum of PD patients can be investigated using positron emission tomography (PET) and the radiotracer *L*-3,4-dihydroxy-6-^18^F-fluorophenylalanine (^18^F-DOPA) (Garnett et al., 1983; Nahmias et al., 1985). ^18^F-DOPA is metabolised in dopaminergic nerve terminals by the enzyme aromatic *L*-amino acid decarboxylase (AADC) to produce ^18^F-DA, which is incorporated into synaptic vesicles and then released into the synaptic cleft following neuronal stimulation (Firnau et al., 1987). In patients with PD, dopaminergic function in the striatum is impaired, and the influx rate constant (*K*_*i*_) of ^18^F-DOPA is significantly decreased in the caudate and putamen (Brooks et al., 1990; Holthoff-Detto et al., 1997; Snow et al., 1993). Dopamine turnover can be measured with extended scanning time, and the effective distribution volume ratio (*EDVR*) of ^18^F-DOPA can be calculated (Sossi et al., 2001). A decreased *EDVR* in the striatum indicates an increase in dopamine turnover, and this was observed in newly diagnosed PD patients relative to healthy controls prior to a decrease in *K*_*i*_ (Sossi et al., 2002). An increase in dopamine turnover was also observed in asymptomatic *LRRK2* mutation carriers before any evidence of changes in *K*_*i*_ (Sossi et al., 2010). This has been hypothesised to be a compensatory mechanism in early PD potentially due to upregulation of AADC decarboxylase activity (Adams et al., 2005; Lee et al., 2000). The rat is easily amenable to PET imaging and *in vivo* dopaminergic compensation has been reported in the 6-hydroxy-dopamine (6-OHDA) lesion rat model of PD (Sossi et al., 2009).

Genetic mouse and rat models of PD have been generated using different approaches with variable success. Although the majority of transgenic and knock-out/knock-in models are in the mouse, there have been several reported rat models. One transgenic rat model has been produced with a human *SNCA* construct with two PD mutations (A30P and A53T) under control of the rat *TH* promoter, and the major deficits reported in this model were olfactory (Lelan et al., 2011). A bacterial artificial chromosome (BAC) transgenic rat model expressing human *SNCA* with the E46K mutation identified a trend for decreased immunostaining for TH in the striatum and α-synuclein aggregates were identified in neuronal processes in the striatum (Cannon et al., 2013). The most representative genetic rat model of PD to date used a human BAC transgene to over-express WT human *SNCA* (BAC-*hSNCA*) (Nuber et al., 2013). Transgenic rats were observed to have significantly decreased expression of TH in the striatum as well as significantly decreased striatal dopamine levels in aged BAC-*hSNCA* rats compared with healthy controls (Nuber et al., 2013). Furthermore, neuritic α-synuclein pathology was identified in the striatum of aged transgenic rats (Nuber et al., 2013).

To date the majority of genetic rat models of PD have been generated by the random insertion of a transgene or the knock-out of a PD-related gene (Creed and Goldberg, 2018). Replicating the precise single amino acid substitution observed in PD kindreds has become possible in the rat with Clustered regularly interspaced short palindromic repeats (CRISPR)/CRISPR-associated protein 9 (Cas9) genome engineering (Yoshimi et al., 2014). Here, we used CRISPR/Cas9 and a donor oligonucleotide in rat zygotes to mutate glycine-51 to aspartic acid (G51D) in rat *SNCA* exon 3 to produce a novel *SNCA*^G51D^ rat model of PD. Striatal dopamine metabolism and turnover were investigated in *SNCA*^G51D/+^ and *SNCA*^G51D/G51D^ rats over the course of ageing using ^18^F-DOPA PET imaging and kinetic modelling.

## MATERIALS AND METHODS

### Point mutation in rat genome with CRISPR/Cas9

CRISPR/Cas9 constructs used were the gRNA_Cloning vector (a gift from George Church, Addgene plasmid # 41824 ; http://n2t.net/addgene:41824 ; RRID:Addgene_41824) and humanised Cas9 nuclease vector (a gift from George Church, Addgene plasmid # 41815 ; http://n2t.net/addgene:41815 ; RRID:Addgene_41815) (Mali et al., 2013). The gRNA sequence was designed, synthesised inserted into the gRNA_Cloning vector into the *Afl*II site. The gRNA sequence is: 5’-GTCGTTCATGGAGTGACAAC-3’. The gRNA and Cas9 vectors were each linearized with *Xho*I and *in vitro* transcribed using a MessageMAXT7 ARCA-Capped Message Transcription Kit (CELLSCRIPT, Madison, WI, USA). A 80-nt ssDNA mutant donor oligonucleotide with the desired 2-bp mutation (G**GA** to G**AT**) was synthesised. The sequence of the ssDNA donor is: 5’-CAATTCTTTTTTTAGGTTCCAAAACTAAGGAGGGAGTCGTTCATGatGTGACAACAGGTAAGCTCTGTTGTCTTTTATCC-3’. The gRNA, *Cas9* mRNA, and ssDNA donor oligonucleotide were co-injected into male pronuclei of F344/Stm rat zygotes. Eleven (11) founder rats were screened by Sanger sequencing. Five (5) founders had mutations in exon 3 of *SNCA*, and one founder had the desired G**GA** to G**AT** (G51D) mutation (Supplementary Fiigure S1). This founder was mated to produce F1 progeny, and the mutation transferred through the germ-line as determined by Sanger sequencing of genomic DNA extracted from ear notches (Figure 1A). Since the mutation introduced a new *Bsp*HI restriction site (Figure 1B), subsequent genotyping was performed by *SNCA* exon 3 PCR (forward primer 5’-TGGTGGCTGTTTGTCTTCTG-3’ and reverse primer 5’-TCCTCTGAAGACAATGGCTTTT-3’), a *Bsp*HI restriction enzyme digest, and agarose gel electrophoresis (Figure 1C).

**Figure 1.**
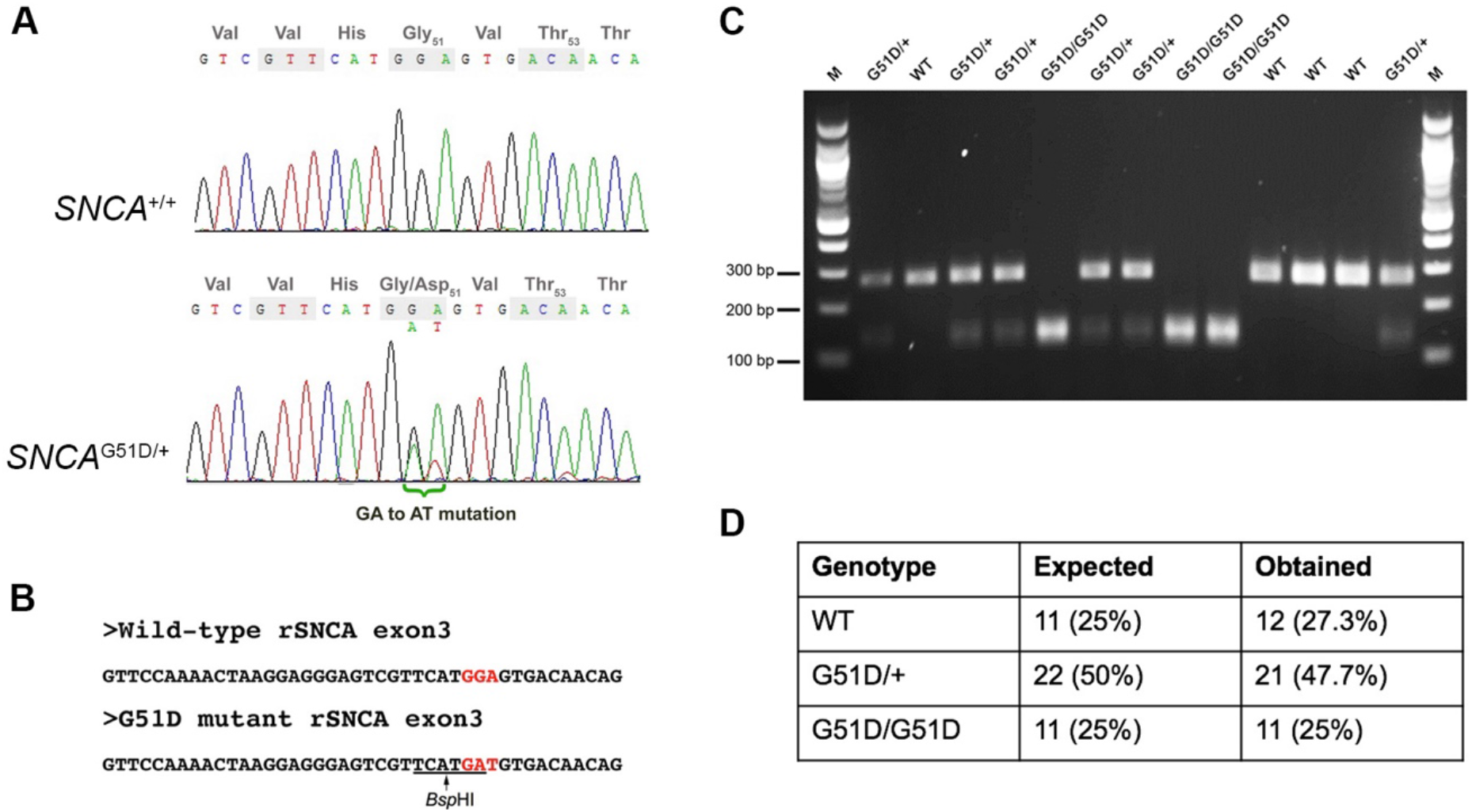
Genotyping of *SNCA*^G51D^ rats. (A) Sanger sequencing of rat *SNCA* exon 3 of a WT and an *SNCA*^G51D/+^ F1 rat revealed a heterozygous mutation (G**<u>GA</u>**/G**<u>AT</u>**) at codon 51. (B) The G51D mutation generated a new *Bsp*HI restriction site (TCATGA) in exon 3. (C) *SNCA* exon 3 PCR and *Bsp*HI restriction enzyme digest can reliably genotype WT, *SNCA*^G51D/+^ and *SNCA*^G51D/G51D^ rats. (D) 44 neonatal (P2, P3) pups from five (5) *SNCA*^G51D/+^ X *SNCA*^G51D/+^ crosses were genotyped, and Mendelian ratios were observed.

### Animal husbandry

The breeding and maintenance of wild-type (WT), *SNCA*^G51D/+^ and *SNCA*^G51D/G51D^ rats, and the *in vivo* experiments conducted were approved by the UK Home Office under project licence PC6C08D7D. WT, *SNCA*^G51D/+^ and *SNCA*^G51D/G51D^ rats were analysed over the course of ageing at 5, 11, and 16 months using ^18^F-DOPA PET imaging (n=3/4 per genotype per age-group).

### PET imaging

^18^F-DOPA was produced using a multi-step nucleophilic fluorination pathway, and used the TRACERlab MX synthesiser and cassette with the nucleophilic precursor to ^18^F-DOPA (ABX Advanced Biochemical Compounds, PEDP-0062-H), with the final product formulated in PBS (Martin et al., 2013).

Imaging was conducted under general anaesthesia and used Isoflurane in 0.6 L/min O_2_ and 0.4 L/min N_2_O_2_. 10 mg/kg Carbidopa (Sigma-Aldrich, C1335), then 10 mg/kg Entacapone (Sigma-Aldrich, SML-0654) were injected intravenously to prevent the peripheral metabolism of ^18^F-DOPA by AADC and catechol-O-methyltransferase (COMT), respectively (Walker et al., 2013a). Thirty minutes after injection of Carbidopa and Entacapone, 18.5 +/- 7.1 MBq (mean +/- SD) of ^18^F-DOPA was injected as a bolus via the tail vein. Activity in the empty syringe was measured, with the activity injected calculated and decay corrected. Dynamic PET imaging lasted 2 hours.

Images were obtained using the nanoScan PET/CT scanner (Mediso Medical Imaging Systems, Budapest, Hungary), with PET/CT data acquired using Nucline™ v2.01 acquisition software. Prior to radiotracer injection a scout view CT image was acquired. Dynamic PET imaging commenced upon injection of ^18^F-DOPA and used a coincidence mode 1-5 and coincidence time window of 5ns. After PET imaging, CT data was acquired (trajectory semi-circular, maximum field of view, 480 projections, 55 kVp and 300 ms, binning 1:4) and used for attenuation correction of PET data.

The PET data that was reconstructed and extended from the olfactory bulbs to the caudal border of the heart. Reconstruction was 3D, dynamic and used the TeraTomo3D reconstruction method. Coincidence mode was 1-3, voxel size 0.4mm x 0.4mm x 0.4mm, used 4 iterations and 6 subsets, and was corrected for scatter, attenuation and randoms. Data was reconstructed into frames comprising 6 frames of 30 sec, 3 frames of 60 sec, 2 frames of 120 sec and 22 frames of 300 sec. All PET imaging data is freely accessible: https://datashare.ed.ac.uk/handle/10283/4014

### Image analysis

Images were analysed using PMOD 3.409 software (PMOD Technologies LLC) and a hand-drawn template. Volumes of interest (VOIs) comprised the left and right striatum, whole striatum, and the cerebellum which was a reference region for non-specific uptake. The same VOI template was used for all rats and were only moved into position over the respective anatomical areas. PET images for display purposes and to aid with VOI drawing were obtained by averaging over 0-120 minutes of emission data then a 1mm x 1mm x 1mm Gaussian filter was applied. Standardized uptake value (SUV) images were produced using rat body weight and the activity injected. Similarly, SUV TACs were calculated as follow SUV (g/ml) = activity concentration in the target VOI (kBq/ml)/(decay corrected amount of ^18^F-DOPA injected (MBq)/weight of the rat (kg)).

Kinetic modelling used the Patlak reference tissue model and Logan reference tissue model to determine the *K*_*i*_ of ^18^F-DOPA in the striatum and the distribution volume ratio (*DVR*) of ^18^F-DOPA in the striatum relative to the cerebellum, respectively (Logan, 2000). The Patlak reference tissue model used 60 min of data and a *t\** of 10 min in accordance with previous studies in Sprague Dawley rats (Kyono et al., 2011). In contrast, the Logan reference tissue model used 120 min of data and *t\** of 30 min, which were parameters optimised from WT F344 rat data and these differ from other methods used to analyse PET data from Sprague Dawley rats (Walker et al., 2013a). The Logan reference tissue model was also used to determine the effective *DVR* (*EDVR*) of ^18^F-DOPA which involved subtracting the TAC (kBq/ml) for the cerebellum from the TACs for the striatum before analysis (Walker et al., 2013a) and used 120 min of data and *t\** of 30 min. *EDVR* can be used to estimate the effective dopamine turnover (Sossi et al., 2002). Differences in *EDVR* between left and right striatum were also investigated by calculating asymmetry in *EDVR* which = (*EDVR* contralateral - *EDVR* ipsilateral)/*EDVR* contralateral (Walker et al., 2013b).

### Statistical analysis

^18^F-DOPA PET data from WT, *SNCA*^G51D/+^, and *SNCA*^G51D/G51D^ rats are presented where appropriate as the mean +/- SEM. Results from age-matched WT, *SNCA*^G51D/+^ and *SNCA*^G51D/G51D^ rats were analysed using a 1-way ANOVA with Tukey’s multiple comparison test.

## RESULTS

### Generation of the *SNCA*^G51D^ rat model

The G51D mutation in PD patients is due to a single nucleotide change (c.152 G>A) that mutates the Gly-51 codon (G**G**T) to G**A**T, encoding for aspartic acid. The rat *SNCA* Gly-51 codon is G**GA**, which requires a 2-nucleotide mutation to produce the Asp-51 codon G**AT**. *Cas9* mRNA, gRNA and ssDNA donor oligonucleotide with GAT codon were co-injected into F344/Stm rat zygotes. Eleven founder rats were screened by Sanger sequencing and 5 F0 rats had mutations in *SNCA*, but only one founder had the desired GGA to GAT (G51D) mutation in exon 3 of *SNCA* (Supplementary Fiigure S1). This F0 founder rat was bred, and F1 animals with the G51D mutation were identified by Sanger sequencing confirming the mutation could transmit through the germ-line (Figure 1A). All subsequent progeny from the sequence-confirmed F1 rats were PCR-genotyped taking advantage of a new *Bsp*HI restriction site introduced by the G51D mutation (Figure 1B,C). Heterozygous *SNCA*^G51D/+^ matings produced progeny in Mendelian ratios suggesting the mutation did not reduce embryonic or fetal viability (Figure 1D).

### Optimisation of ^18^F-DOPA PET imaging in Fischer 344 rats

The methods used for the reconstruction of *in vivo* PET imaging experiments were optimised since the rat striatum is a small structure and the optimal acquisition and reconstruction of PET data is scanner dependent. A homogenous solution of ^18^F-FDG was used in a National Electrical Manufacturers Association (NEMA) NU-4 mouse image quality (IQ) phantom (Harteveld et al., 2011). Imaging data was acquired and reconstructed using variations of several parameters (Supplementary Fiigure S2). The most promising reconstruction scenario used iterative methods and employed 4 iterations and 6 subsets (4i6S) at a normal resolution and a coincidence mode of 1-3. The *in vivo* imaging and kinetic modelling methods were optimised using data obtained from preliminary experiments using two WT F344 rats (Figure 2). Time activity curves (TACs) and standardized uptake values (SUV) were calculated and plotted for striatum and cerebellum (Figure 2B,C). The cerebellum was used as a reference region since it lacks AADC expression (Baker et al., 1991). Plotting the SUV ratio (SUVr) of the striatum and cerebellum for both rats showed the data was consistent and that the activity reached a pseudo-equilibrium between 50 min and 85 min (Figure 2D). This data indicated that methods used for kinetic modelling in recent PET imaging studies of models of PD were optimal (Kyono et al., 2011). Kinetic modeling using Patlak and Logan graphical analysis used data collected over 60 min and 180 min, respectively (Figure 3)

**Figure 2.**
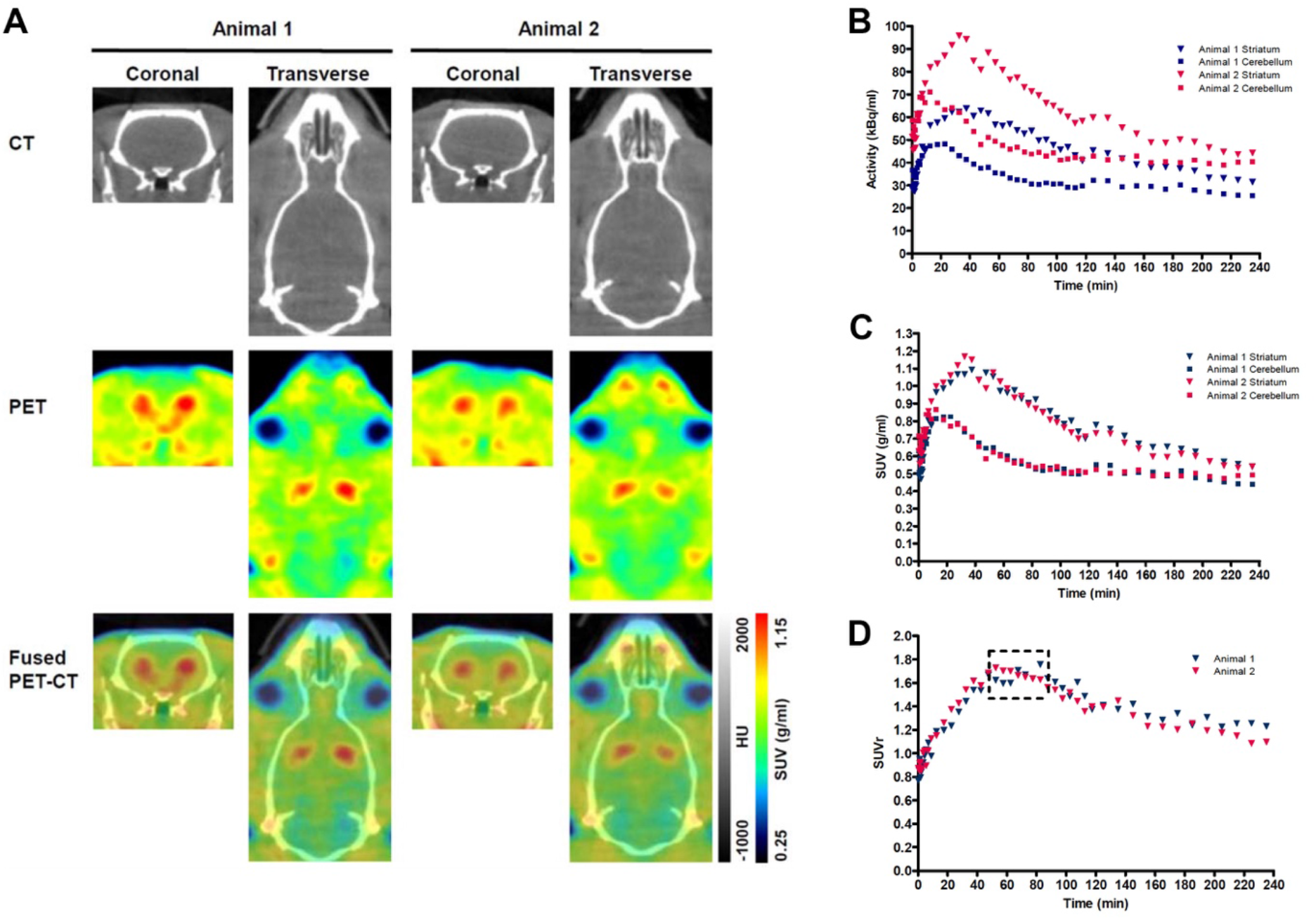
Representative PET-CT images and Standardized uptake value (SUV) time activity curves (TACs) for wild-type (WT) rats. (A) PET-CT images of two WT rats are shown in coronal and transverse planes. ^18^F-DOPA PET data averaged over frames 1-33 and smoothed using a 1mm x 1mm x 1mm Gaussian filter. HU-Hounsfield Units. Specific activity (B), SUV TAC data (C), and SUV ratio (SUVr) data (D) for the specific uptake of ^18^F-DOPA into the striatum relative to the cerebellum is shown for two WT rats. The dashed box indicates the phase of pseudo-equilibrium (50 - 85 min).

**Figure 3.**
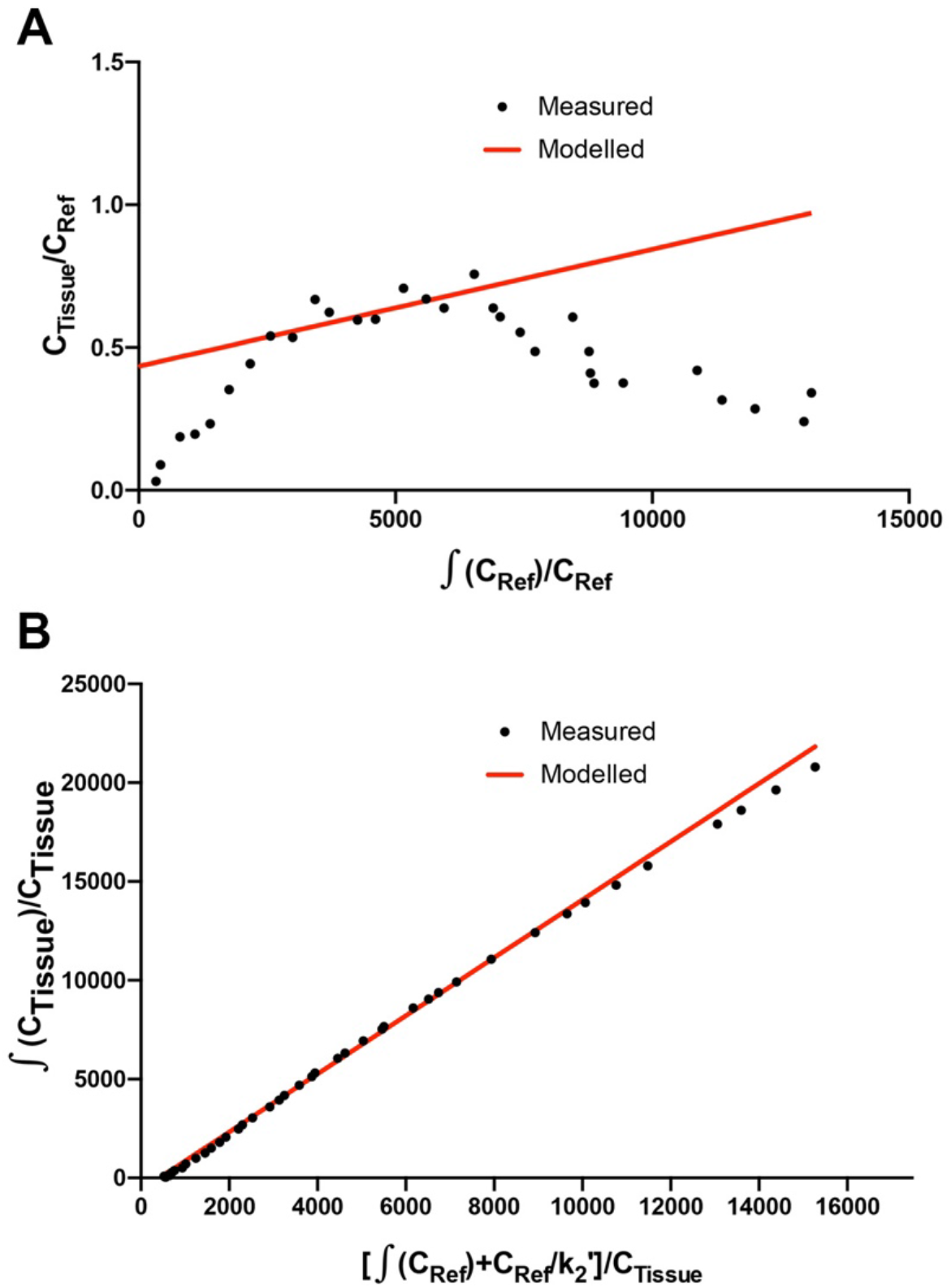
Patlak and Logan analysis for WT rats. (A) Patlak graphical analysis of 60 min of data from the whole striatum of a WT rat relative to cerebellum. (B) Logan graphical analysis of 180 min of data from the whole striatum of a WT rat.

### ^18^F-DOPA PET imaging reveals a deficit in dopamine turnover

WT, *SNCA*^G51D/+^ and *SNCA*^G51D/G51D^ rats were subjected to ^18^F-DOPA PET at 5, 11, and 16 months of age using the conditions optimised above. Data was analysed to determine SUV TACs from WT, *SNCA*^G51D/+^ and *SNCA*^G51D/G51D^ rats (Figures 4A), and plotted to show the specific uptake of ^18^F-DOPA into the striatum compared with the cerebellum.

**Figure 4.**
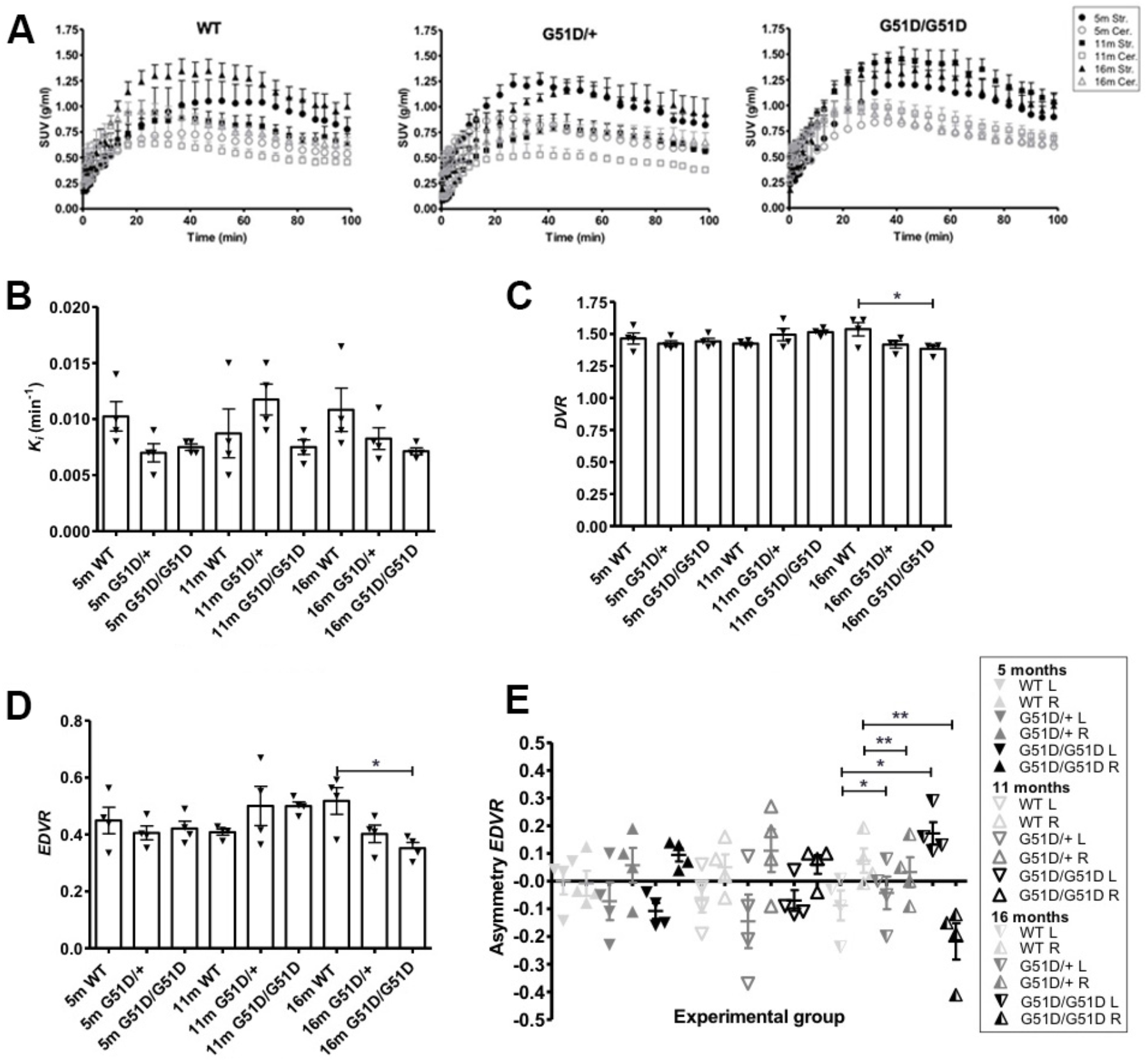
Kinetic modelling of ^18^F-DOPA PET data from WT, *SNCA*^G51D/+^ and *SNCA*^G51D/G51D^ rats at 5, 11 and 16 months of age. (A) Standardized uptake value (SUV) time activity curves (TACs) for all rat genotypes – WT, *SNCA*^G51D/+^, and *SNCA*^G51D/G51D^ rats, n=4 per genotype per age-group. m-months old, Str.-Striatum, Cer.-Cerebellum. (B) The mean *K*_*i*_ in the striatum of 5-16 month old WT, *SNCA*^G51D/+^ and *SNCA*^G51D/G51D^ rats. (C) There was a significant decrease in mean *DVR* of ^18^F-DOPA in 16 month old *SNCA*^G51D/G51D^ rats when compared with age-matched WT rats (p=0.0348), and (D) also a significant decrease in *EDVR* of ^18^F-DOPA in 16 months *SNCA*^G51D/G51D^ rats compared with age-matched WT rats (p=0.0215). (E) Mean left-right asymmetry in *EDVR* was greatest in 16 month old *SNCA*^G51D/+^ and *SNCA*^G51D/G51D^ rats. Data shows the mean, SEM and individual data-points. n=4 per genotype per age-group. One-way ANOVA with Tukey’s multiple comparison test. L-left striatum, R-right striatum, m-months old.

The mean *K*_*i*_ values of ^18^F-DOPA in 5-16 month old *SNCA*^G51D/+^ and *SNCA*^G51D/G51D^ rats compared with age-matched WT rats were not significantly different (Figure 4B). The mean *DVR* and *EDVR* of ^18^F-DOPA in the striatum relative to the cerebellum was significantly decreased in 16 month old rats compared to WT rats, but these differences were not observed at 5 and 11 months of age (Figure 4C,D). The *EDVR* of ^18^F-DOPA is the ratio of the distribution volumes of ^18^F-DOPA in the specific and precursor compartments reduced by the factor *k*_*2*_/(*k*_*2*_ + *k*_*3*_), and since *EDVR* is estimated to be the inverse of effective dopamine turnover (Sossi et al., 2002), the results indicate an increase in mean dopamine turnover in 16 month *SNCA*^G51D/G51D^ rats compared with age-matched WT rats. Interestingly, the mean asymmetry in the *EDVR* of ^18^F-DOPA in the left and right striatum of 16 month old *SNCA*^G51D/G51D^ and *SNCA*^G51D/+^ rats were significantly different than age-matched WT rats (Figure 4E). The *EDVR* asymmetry in *SNCA*^G51D/G51D^ rats was between -0.4 and 0.3, while the normal range determined for Sprague Dawley rats is between -0.1 and 1.0 (Walker et al., 2013b).

## DISCUSSION

The aim of the study was to characterise a novel *SNCA*^G51D^ rat model of Parkinson’s disease using ^18^F-DOPA PET imaging. Experiments were conducted over an ageing time-course, since phenotypes were anticipated to worsen with time. In patients with PD, striatal dopaminergic function decreases prior to degeneration of nerve terminals (Morrish et al., 1998; 1995; Nurmi et al., 2001).

^18^F-DOPA PET imaging data indicated no significant differences in mean *K*_*i*_ of ^18^F-DOPA in the striatum of ageing *SNCA*^G51D/+^ and *SNCA*^G51D/G51D^ rats compared with age-matched WT rats (Figure 4B). However, there was a significant decreased mean *DVR* and *EDVR* of ^18^F-DOPA in the striatum of 16 month old *SNCA*^G51D/G51D^ rats compared with age-matched WT rats (Figure 4C,D), which indicates increased dopamine turnover in the aged *SNCA*^G51D/G51D^ rats.

In early PD the *EDVR* of ^18^F-DOPA in the striatum relative to the cerebellum is significantly decreased compared with healthy controls, which indicates increased effective dopamine turnover and this is likely due to a compensatory change in response to striatal dysfunction (Sossi et al., 2002). This data is supported by results from ^18^F-DOPA PET imaging studies of asymptomatic MPTP lesioned monkeys which have also implicated increased effective dopamine turnover as a compensatory mechanism (Doudet et al., 1998). The significantly decreased mean *EDVR* in 16 month old *SNCA*^G51D/G51D^ rats may indicate a compensatory change in dopaminergic function in the striatum of *SNCA*^G51D/G51D^ rats following putative striatal dysregulation.

In PD the *K*_*i*_ of ^18^F-DOPA in the striatum is significantly decreased compared with healthy controls (Brooks et al., 1990; Burn et al., 1994; Holthoff-Detto et al., 1997; Rinne et al., 2000), with *K*_*i*_ in the putamen reaching 57-80% of normal levels before symptoms of PD develop (Morrish et al., 1995; 1998; Nurmi et al., 2001). The lack of change in mean *K*_*i*_ in the striatum of *SNCA*^G51D/+^ and *SNCA*^G51D/G51D^ rats suggests any potential phenotypes are being sufficiently compensated for in this model and reflects what is observed in patients with prodromal PD. In chemical lesion models of PD, TH has been found to be significantly increased in the striatum post lesioning, a compensatory mechanism in response to nerve damage, which either results in increased TH protein expression or involves morphological changes such as the expansion of nerve terminals (Bezard and Gross, 1998; Blanchard et al., 1995). Compensatory changes in dopaminergic terminals have also been shown to involve the upregulation of AADC activity and the downregulation of DATs (Adams et al., 2005; Lee et al., 2000). In 6-OHDA lesion models of PD, PET imaging studies have identified a complex relationship between the *k*_*ref*_ of ^18^F-DOPA and the binding potential of ^11^C-DTBZ (denervation severity) (Walker et al., 2013a).

Results measuring asymmetry in the *EDVR* of ^18^F-DOPA showed decreased mean *EDVR* in the left striatum when compared to the right in 5 out of 6 groups of rats, with greatest asymmetry in 16 month old *SNCA*^G51D/G51D^ rats (Figure 4E). Reasons that may explain the predilection for the left side could be due to the unique anatomy of the blood vessel supply to the brain in F344 rats which may result in non-uniform perfusion in this strain of rat (Iwasaki et al., 1995).

*SNCA*^G51D^ rats were generated using CRISPR/Cas9 technology to model the G51D α-synuclein mutation in humans. The G51D model has good construct validity, and mutant α-synuclein is expressed from the endogenous rat locus. *SNCA*^G51D^ rats have the potential to model the widespread neurological abnormalities found in PD and may be a more representative model of PD than the focal 6-OHDA lesion model, which has previously been studied using ^18^F-DOPA PET imaging (Kyono et al., 2011; Walker et al., 2013a). Since rodent models of human disease are almost always less severe, we expected the *SNCA*^G51D^ rats to have phenotypes similar to early stages of the disease (Petrucci et al., 2016). *SNCA*^G51D^ may exhibit a subtle phenotype in rats due to protective factors such as β-synuclein which have been shown to the ameliorate PD-like phenotypes in mice (Fan et al., 2006; Hashimoto et al., 2001). Furthermore, rodents may lack key triggers or cellular components necessary for exhibiting a full PD phenotype. Differences in the *SNCA* sequence between rodents and humans may explain some of the difficulties in modelling genetic PD, since rodents, indeed most vertebrates, encode a threonine at position 53 which is a cause of PD in humans, and human A53T mice have demonstrated exaggerated motor deficits and α-synuclein pathology following removal of endogenous mouse α-synuclein (Cabin et al., 2005).

## Conclusion

*SNCA*^G51D/G51D^ rats show a significant increase in dopamine turnover in the striatum at 16 months of age, but not a significant decrease in Ki – dopamine synthesis and storage. These findings mimic the early stages of Parkinson’s, and may reflect the compensatory changes in the dopaminergic system observed in humans. *SNCA*^G51D^ rats represent an interesting model of early PD pathophysiology, and provide a tractable platform for investigating additional genetic or environmental triggers of Parkinson’s.

## Acknowledgements

This work was funded by a Carnegie Trust PhD Scholarship to VM, Parkinson’s UK Senior Fellowship to TK, a Wellcome Trust ISSF award to TK and AT. We thank Ms Yayoi Kunihiro for assistance in generation of the *SNCA*^G51D^ rat model.

## Competing interests

The authors declare no competing interests.

## Authors’ contribution

VM, AT, and TK devised the study. The *SNCA*^G51D^ rat model was generated by TM (Kyoto University). KSD (University of Edinburgh) established genotyping of *SNCA*^G51D^ rats, maintained the rat colony, and conducted preliminary studies. The ^18^F-DOPA radiotracer was synthesised by TW and CL (University of Edinburgh), and CCA (University of Edinburgh) provided assistance during *in vivo* ^18^F-DOPA PET imaging experiments. VM and AT designed and conducted PET experiments and data analysis. VM and TK wrote the manuscript and it was edited by AT.

## Notes

https://datashare.ed.ac.uk/handle/10283/4014

### Competing Interest Statement

The authors have declared no competing interest.

