## Supplementary Information for "*In vivo* ^18^F-DOPA PET imaging identifies a dopaminergic deficit in a rat model with a G51D α-synuclein mutation"

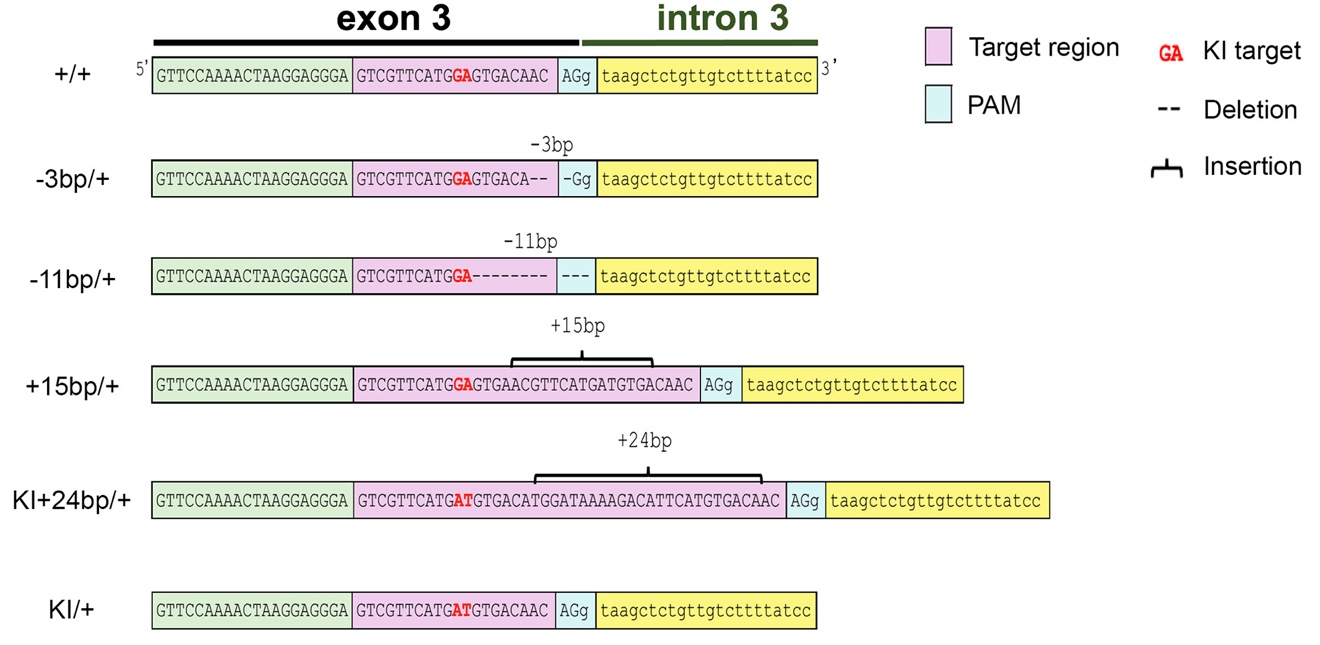

**Supplementary Figure S1. DNA sequence of CRISPR/Cas9 founder rats.** Five (5) of 11 F0 rats had mutations in the rat *SNCA* locus. Only one founder animal had the correct knock-in (KI) target with the GA to AT mutation without additional mutations.

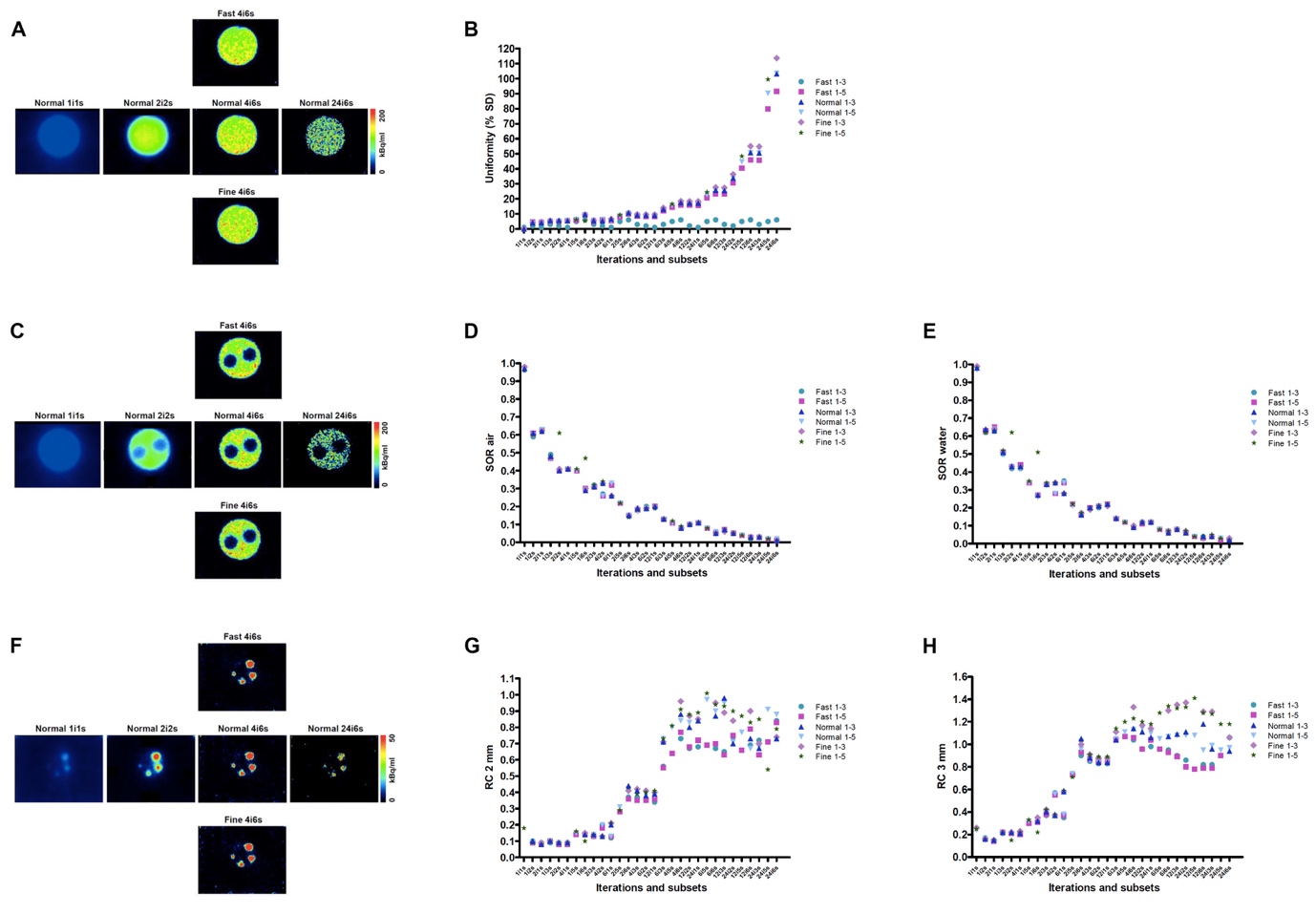

**Supplementary Figure S2. PET IQ Phantom data.** The phantom was filled

with 3.8 MBq of ^18^F-FDG solution. (A) Images of the central chamber after reconstructing the PET data using different methods varying the resolution (fast/normal/fine) and the number of iterations and subsets used for reconstruction. (B) Effect of the reconstruction method on the % standard deviation (% SD) in image uniformity. (C) Images of air- and water-filled inserts of the phantom. (D,E) Effect of spillover ratio (SOR) of activity into air (D) and water (E). (F) Images of rods of differing diameters (1mm – 5mm) in the phantom. (G,H) Recovery coefficients (RC) for 2 mm rods (G), and 3 mm rods (H).
